# Temporal profiling of *Salmonella* transcriptional dynamics during macrophage infection using a comprehensive reporter library

**DOI:** 10.1101/2023.09.27.559620

**Authors:** Taylor H. Nguyen, Oscar R. Diaz, Manohary Rajendram, Daniel S.C. Butler, Benjamin X. Wang, Jay C. D. Hinton, Denise Monack, Kerwyn Casey Huang

## Abstract

The transcriptome of *Salmonella enterica* serovar Typhimurium (*S*. Tm) dynamically responds to the rapid environmental shifts intrinsic to *S.* Tm lifestyle, exemplified by entry into the *Salmonella*-containing vacuole (SCV) within macrophages. Intracellular *S*. Tm must respond to the acidity of the SCV, accumulation of reactive oxygen/nitrogen species, and fluctuations in nutrient availability. Despite thorough RNA-seq-based investigations, the precise transcriptional timing of the expression of many secretion systems, metabolic pathways, and virulence effectors involved in infection has yet to be elucidated. Here, we construct a comprehensive library of GFP-reporter strains representing ∼3,000 computationally identified *S.* Tm promoter regions to study the dynamics of transcriptional regulation. We quantified promoter activity during *in vitro* growth in defined and complex media and throughout the timeline of intracellular infection of RAW 246.7 macrophages. Using bulk measurements and single-cell imaging, we uncovered condition-specific transcriptional regulation and population-level heterogeneity in the activity of virulence-related promoters, including SPI2 genes such as *ssaR* and *ssaG*. We discovered previously unidentified transcriptional activity from 234 genes, including ones with novel activity during infection that are associated with pathogenecity islands and are involved in metabolism and metal homeostasis. Our library and data sets should provide powerful resources for systems-level interrogation of *Salmonella* transcriptional dynamics.

## Introduction

*Salmonella* serovars are responsible for human diseases ranging from gastroenteritis to systemic infections. *Salmonella enterica* serovar Typhi only infects humans and is responsible for typhoid fever, while *Salmonella enterica* serovar Typhimurium (*S*. Tm) has a broad host range and can survive in the wider environment. *S*. Tm infections usually result in self-limiting gastroenteritis in humans and systemic typhoid-like disease in mice^1^. *Salmonella* infections, which account for ∼50% of foodborne illnesses worldwide^2^, typically result from exposure to contaminated food or water. The emergence of multidrug-resistant *Salmonella* strains is now posing a major global health risk^3–6^. During systemic *Salmonella* infections, the pathogen penetrates the gut epithelial barrier and preferentially infects phagocytes within the lamina propria^7^. The ability of *Salmonella* to proliferate within macrophages in the *<u>S</u>almonella*-<u>c</u>ontaining <u>v</u>acuole (SCV) is a hallmark of systemic disease^8^.

*S*. Tm can invade macrophages via upregulation of *Salmonella* pathogenicity island 1 (SPI1), which encodes the type 3 secretion system 1 (T3SS-1), associated virulence effectors, and regulators^9^. Following entry into macrophages, *S*. Tm cells respond to the nutrient-restricted and acidified environment of the SCV through transcriptional upregulation of *Salmonella* pathogenicity island 2 (SPI2) and expression of the type 3 secretion system 2 (T3SS-2), which leads to injection of *S*. Tm-derived virulence factors^10^. These virulence factors contribute to a variety of outcomes including skewing of macrophage polarization state, formation of *Salmonella*-induced filaments (elongated tubes that protrude from the SCV to enhance nutrient acquisition), modification of the SCV environment to minimize bacterial clearance, and inhibition of antibacterial pathways within the host cell^10^.

Quantification of the *S*. Tm transcriptome using RNA-seq revealed patterns of gene regulation under a variety of *in vitro* conditions and during macrophage infection^11–13^. Previous studies suggested that *S.* Tm responds to the macrophage intracellular environment by shifting from initial expression of genes within SPI1 to subsequent expression of SPI2 genes in the first 8 h of infection, coinciding with a transcriptional shift of the infected host macrophage^14^. An outstanding challenge is to understand the physiological adaptations and transcriptional dynamics that occur in *S.* Tm throughout intracellular replication, especially during host metabolic reprogramming and nutrient sequestration^15^.

Through an intricate regulatory network, *Salmonella* can adapt its transcriptional program within 4 min upon encountering a new environment^16^. Given the short half-life of mRNA transcripts^17^, capturing response dynamics with a high temporal resolution is vital. Furthermore, the identification of additional virulence-related genes and determination of their expression dynamics during infection remain important challenges, and a deeper understanding of the transcriptional response of *Salmonella* to various environmental perturbations should empower development of antimicrobial treatments to *Salmonella* infection. Technical limitations and the expense of RNA-seq are obstacles to measurement of the genome-wide transcriptional profile with high temporal resolution. An alternative transcriptional profiling approach involves engineering strains in which a promoter is fused to a reporter, such as a fluorescent protein, to enable measurement of promoter activity over time. This approach has been facilitated by the development of highly stable, fast-folding GFP variants that avoid delays in activity read-out^18^. A comprehensive library of such promoter reporters was constructed and used to quantify the dynamics of the *Escherichia coli* transcriptome, enabling the discovery of transcriptional regulatory regions^19,20^. Previous studies using *S.* Tm promoter reporters were targeted to specific pathways of interest such as the response of biofilm-associated promoters to biofilm inhibitors, of metabolic genes upon entry into the SCV, and stress responses during intracellular infection^21–23^; a comprehensive characterization of the dynamics of the *S.* Tm transcriptome has not been conducted.

Here, we report the construction of a comprehensive reporter library containing ∼3,000 promoter regions identified from the *S.* Tm genome fused to GFP and its application to measuring the dynamics of *S.* Tm transcriptional regulation. We profiled changes in *S*. Tm promoter activity across defined and complex media conditions, capturing condition-specific regulation of promoters and the global response to *in vitro* conditions that mimic aspects of the intra-macrophage environment. Using fluorescence microscopy, we demonstrate that our library can be used to quantify heterogeneity in promoter activity across a population of single bacterial cells. To determine the transcriptional response of *S*. Tm to the intracellular macrophage environment, we used time-lapse fluorescence microscopy of RAW 264.7 macrophage-like cells infected with each reporter strain in the library individually. Our experimental screens provided the first evidence of activity for 234 promoters. We found that the *mntS* promoter region is dependent on environmental manganese concentrations and is active during macrophage infection, highlighting the importance of metal homeostasis during pathogenesis. By capturing transcriptional dynamics, we identified a metabolic shift in *S.* Tm to the Entner-Doudoroff (ED) pathway during the later stages of intracellular infection. In total, our library and analyses uncovered temporal patterns of transcriptional regulation involving *S*. Tm genes related to metabolism, metal acquisition, and pathogenicity.

## Results

### Construction of a comprehensive library of *Salmonella* Typhimurium transcriptional reporters

Our goal was to construct a comprehensive arrayed library of GFP reporter fusions to every lead-operon (primary) promoter in *S.* Tm. To this end, we identified 2,901 promoters using an unbiased computational approach. First, we extracted the intergenic distances between NCBI-annotated coding sequences (CDS) across the *S.* Tm strain SL1344 genome. Genes with intergenic distances <40 bp were considered to be transcribed as part of an operon and were excluded from further analysis. For the remaining 2,901 CDS, we defined the promoter region as the 350 bp upstream of and including the translational start site (TSS, start codon). Of these primary promoters, the transcriptional start sites of 1,850 were experimentally validated in previous studies that enriched for *S.* Typhimurium 4/74 primary transcripts via differential RNA-seq (dRNA-seq)^13^. The remaining 1,051 were considered putative. We did not consider potential promoters within genes or regulation that occurs from sequences after the start codon. We did not include regions downstream of the TSS because we reasoned that including the N-terminal signal sequence that translocates proteins to the periplasmic space and the inner/outer membrane would interfere with the folding and stability of our GFP fusion, a hypothesis that was borne out for a small set of randomly selected membrane proteins (**Fig. S1**). The final reporter constructs in our library, which contain the promoter regions followed by the *mGFPmut2* sequence^18^, were cloned into a common plasmid backbone that includes the pSC101 origin of replication, the *mob* mobilization region to enable conjugative transfer between strains, and a chloramphenicol-resistance (*Cm^R^*) cassette (**Fig. 1A**).

**Figure 1:**
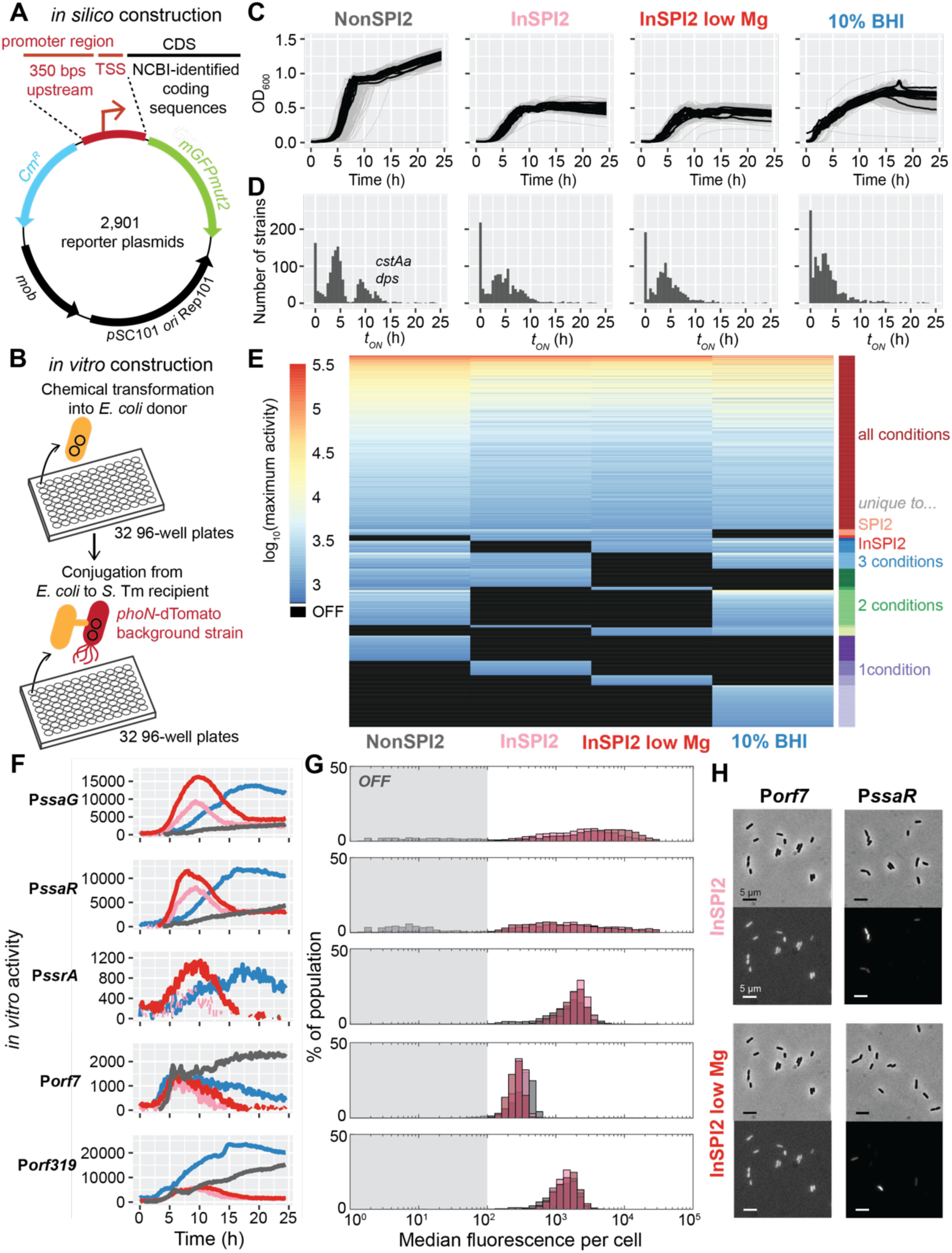
High-throughput construction of a comprehensive *S*. Tm reporter library enables activity profiling across *in vitro* media conditions. A) Plasmid map of the reporter-GFP fusions in the library. Promoter regions were identified as the 350 bp upstream of and including the TSS, followed by the *mGFPmut2* sequence, the pSC101 origin of replication, the *mob* mobilization region for conjugative transfer, and a chloramphenicol (Cm)-resistance (*Cm^R^*) cassette. B) High-throughput cloning strategy used to generate the library. Plasmids were chemically transformed into *E. coli*, and positive transformants were used for conjugation into a *S.* Tm parental strain with *dTomato* integrated at the *phoN* locus. Positive *E. coli* and *S.* Tm transformants are stored in 32 96-well plates. C) Background-subtracted optical density (OD_600_) dynamics for all reporter strains in four *in vitro* media conditions (NonSPI2, InSPI2, InSPI2 low Mg, and 10% BHI). Growth curves for reporter strains are shown in gray, and background (promoter-less) strains are shown as thick black lines. D) Histograms of the time *t*_ON_ at which reporters first exhibited significant activity in each medium (NonSPI2, InSPI2, InSPI2 low Mg, and 10% BHI). E) Maximum activity for the 1,850 reporters that turned ON in at least one *in vitro* condition. Activity is reported as parent-subtracted GFP normalized by background-subtracted OD_600_. Reporters are ordered by their shared activity across media. Black values denote OFF activity. F) Promoter activity dynamics for five SPI2 operon reporters (*ssaG*, *ssaR*, *ssrA*, *orf7*, *orf319*) in NonSPI2 (grey), InSPI2 (pink), InSPI2 low Mg (red), and 10% BHI (blue). Activity is reported as GFP (parent-subtracted to account for autofluorescence) normalized to blanked OD_600_ (to account for changes in cell number). A promoter was defined as ON based on comparison to a dynamic estimate of background noise (**Methods**). ON activity is denoted by solid lines, whereas OFF activity is denoted by dashed lines. G) Histograms of median GFP intensity for >1,000 cells in NonSPI2 (grey), InSPI2 (pink), and InSPI2 Low Mg (red) for the strains in (F). P*ssaG* and P*ssaR* cells exhibited substantial fluorescence heterogeneity. Strains were imaged after 8 h of growth in each medium. H) Representative phase (left) and GFP (right) images of P*orf7* and P*ssaR* cells after 8 h of growth in InSPI2 and InSPI2 low Mg. Fluorescence contrast was adjusted for each strain to highlight population heterogeneity. Scale bar: 5 µm.

Plasmids were assembled, sequence verified, and arrayed into 96-well plates. Using a high-throughput cloning strategy in 96-well plates (**Methods**), the arrayed plasmid library was transformed into chemically competent *E. coli* NT11164 (**Table S1**), a strain that requires diaminopimelic acid for growth and contains an integrated high-frequency recombination plasmid for conjugation into *S.* Tm^24^ (**Fig. 1B**, **Methods**). The plasmids were then conjugated into a *S.* Tm SL1344 strain containing the *dTomato* gene (which encodes the fluorescent protein dTomato) at the *phoN* locus, an established reporter that is up-regulated by >50-fold during intracellular macrophage infection^13,25^. This *S.* Tm SL1344::*phoN-dTomato* strain retains comparable infectious capacity to wild-type *S.* Tm *in vivo*^26,27^. Library construction resulted in successful cloning of 99.1% (2,874 of 2,901) of the reporter plasmids into *S*. Tm. We sequence verified the promoter region of a randomly selected subset of ∼150 strains including ones that were the focus of follow-up experiments (**Table S2**); the expected identify was confirmed in almost all cases, and mismatches attributed to human error during colony picking were not used for downstream analyses. Our *E. coli* collection of *S.* Tm reporter plasmids is stored in 96-well format to simplify future conjugation into other strains of interest, and the final arrayed library of GFP reporter fusions in *S.* Tm SL1344::*phoN-dTomato* enables the interrogation of *S.* Tm transcriptional dynamics and regulation using high-throughput screens in a fluorescence plate reader or via microscopy, as we demonstrate below.

### Profiling *S.* Tm transcriptional dynamics across *in vitro* conditions

To profile infection-relevant transcriptional programming, we characterized our library during *in vitro* growth in media that represent the key physiochemical aspects of the intracellular macrophage environment. Previous work probing the *Salmonella* transcriptional response used a defined minimal medium to replicate the low pH, low phosphate (P_i_), low magnesium (Mg^2+^), and low nutrient features of the SCV, an intracellular environment within macrophages that induces expression of SPI2^12,28^. Hence, we screened the library in the following defined media: NonSPI2 (pH 7.4, 25 mM P_i_, 1 mM MgSO4), InSPI2 (pH 5.8, 0.4 mM P_i_, 1 mM MgSO4), and InSPI2 low Mg (pH 5.8, 0.4 mM P_i_, 10 µM MgSO_4_). We also screened the library in 10% Brain Heart Infusion (BHI) to probe transcription in a gut-relevant setting^29^, hypothesizing that this rich (undefined) medium would reveal distinct transcriptional programming. Measurements of reporter activity were carried out in a fluorescence plate reader in 384-well black-walled plates by measuring OD_600_ and GFP every ∼10 min over 24 h to generate an *in vitro* data set containing >3.3 million data points.

Virtually all strains in the promoter library exhibited growth kinetics similar to wild-type *S.* Tm (**Fig. 1C**), indicating that the presence of the reporter plasmid does not affect fitness. We quantified the activity of each promoter (hereafter referred to for a given gene as P*gene*) based on the background-subtracted GFP signal (difference between the fluorescence of the strain of interest and the autofluorescence of the parent strain without a plasmid), normalized by background-subtracted OD_600_ to account for changes in cell density. A promoter was defined as “ON” based on comparison to a dynamic estimate of background noise (**Fig. S2**, **Methods**).

The time at which promoter activity was first considered ON (*t*_ON_) was broadly distributed across library strains and conditions (**Fig. 1D**), suggesting that the library can probe transcriptional dynamics in all phases of growth from lag to stationary. In NonSPI2, *t*_ON_ exhibited a bimodal distribution, with the second peak consisting of reporters for genes related to stationary phase such as *dps* (*t*_ON_=8.8 h) and *cstAa* (*t*_ON_=11.9 h), which encode proteins responsible for DNA protection in stationary phase^30^ and escape from carbon starvation^31^, respectively. 64.4% (1,850 of 2,874) of the strains in our library were ON in at least one *in vitro* condition. 50.1% (1,440) of the strains turned ON in NonSPI2, 41% (1,177) in InSPI2, 37.9% (1,089) in InSPI2 low Mg, and 50.6% (1,454) in 10% BHI. Based on the maximum activity across all time points in each of the four conditions, promoters clustered into condition-specific profiles (**Fig. 1E**).

To evaluate the efficacy of our library, we compiled a list of genes related to SP1, T3SS-1, SPI2, and T3SS-2 (**Table S3**) and quantified their activity in the four *in vitro* conditions. Promoters from the SPI2 pathogenicity island (e.g., *orf7*, *orf319*, *ssaG*, *ssaR*, *ssrA*) (**Fig. 1F**), regulators (e.g., *phoP*, *hilA*, *ompR*, *sirC*), structural proteins (e.g., *ompC*, *ompF*, *sicA*), effectors (e.g., *cigR*, *gtgE*, *pipB*, *pipB2*, *sifB*, *sopD2*, *sopE*, *sseJ*, *sseK3*, *sspH2*, *steA*), and other miscellaneous proteins (e.g., *phoN*, *pagC*, *pagO*, *pagD*, *pagK*) that are canonically up-regulated during infection showed high activity (**Fig. S3**), particularly in the InSPI2 conditions (high or low Mg)^12,28,32^. However, contrary to expectations from RNA-seq measurements^12^, some reporters (*ssaB*, *ssaM*, *sscB*) did not exhibit activity in InSPI2 media, indicating that these individual constructs may be missing key regulatory sites. Thus, for downstream analyses, we focused on the reporters with positive signal rather than potential false negatives.

In addition to promoter activity dynamics, the strains in our library enable quantification of population-level heterogeneity in gene expression (e.g., via flow cytometry or single-cell imaging). Previous work showed that some *Salmonella* virulence factors are expressed heterogeneously across single cells, and this phenotypic variation can play an integral role in *Salmonella* virulence^33^. For example, upon SCV invasion, a subpopulation of *Salmonella* cells transition into a non-replicating persister state that can better withstand long-term host-induced damage and antibiotic treatment^34^. Hence, we sought to quantify population-level heterogeneity for five SPI2-operon promoters that turned ON in InSPI2 conditions. We collected fluorescence images of >1,000 cells for each strain after 8 h of growth (approximately when these promoters reached maximum activity in InSPI2) in NonSPI2, InSPI2, and InSPI2 low Mg and quantified the distribution of GFP fluorescence intensity across the population (**Fig. 1G**). These strains displayed a wide range of population-level behaviors. P*orf7* cells exhibited a relatively narrow distribution of GFP fluorescence intensities, whereas P*ssaR* cells exhibited a wide range of intensities that spanned more than two orders of magnitude (**Fig. 1H**). These data also confirmed that the P*orf7* reporter, whose bulk signal was only slightly above our background noise threshold, indeed turned ON at the single cell level, providing confidence for the positive hits from our bulk screen that were near the background threshold. These results demonstrate that our library and experimental setup produces the expected induction of several genes in the tightly regulated SPI2 pathogenicity island, and that the reporters enable quantification of *in vitro* transcriptional dynamics through bulk and single-cell measurements.

### Distinct *Salmonella* promoter activity dynamics in infection-relevant media

In the four growth conditions, the reporter strain library exhibited a wide range of maximum promoter activities and of times to reach maximum activity (*t*_max_) (**Fig. 2A**). Maximum activity was not correlated with *t*_max_, emphasizing the importance of quantifying the full range of dynamic behaviors to comprehensively profile *S.* Tm transcriptional activity. We focused on the 895 promoters that turned on in all three SPI2 conditions (NonSPI2, InSPI2, InSPI2 low Mg) (**Fig. 2B**) and compared activity normalized to its maximum across reporters. Clustering of normalized promoter activity in NonSPI2 revealed sets of promoters with qualitatively distinct patterns (e.g., constitutive, late, pulsatile) (**Fig. 2C**). Ordering the strains by *t*_max_ revealed that promoters collectively reached maximum activity across the entire 24 h in NonSPI2 (**Fig. 2D, left**). To compare dynamic behaviors across conditions, the same analyses were performed for promoter activity in InSPI2 and InSPI2 low Mg (**Fig. 2D**). In contrast to NonSPI2, *t*_max_ values in InSPI2 and InSPI2 low Mg were much more narrowly distributed. In these two conditions, reporter responses resembled a pulse in which activity decreased dramatically after reaching a maximum at ∼10 h (**Fig. 2D, middle and right**), with *t*_max_ for many promoters shifted to an earlier stage of growth compared with NonSPI2. Thus, *S.* Tm displays distinct transcriptional dynamics in SPI2-inducing versus non-inducing conditions.

**Figure 2:**
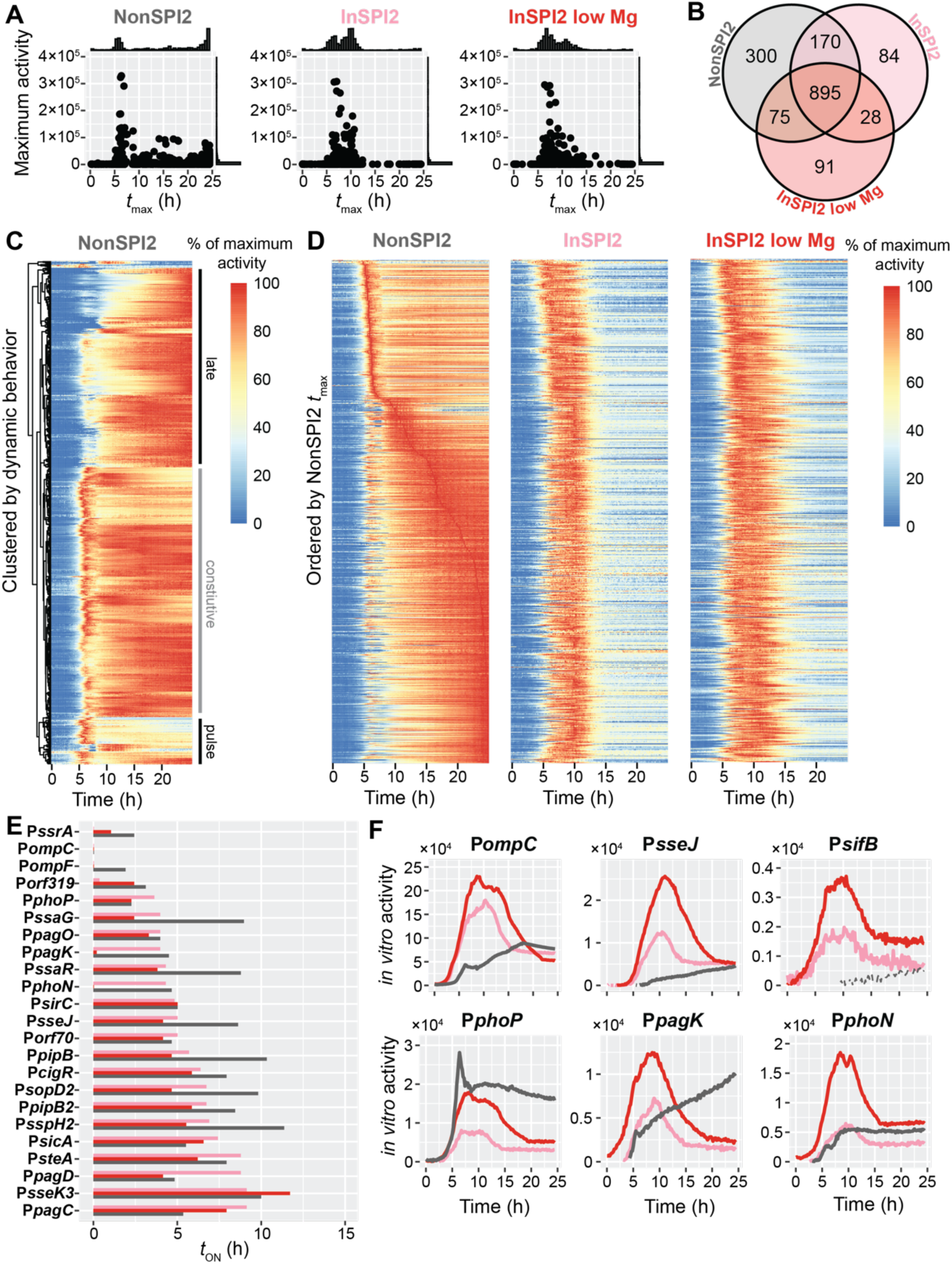
Reporter library reveals system-level shifts in promoter activity dynamics in macrophage-mimicking media. A) Comparisons of time to maximum activity (*t*_max_) and maximum promoter activity in NonSPI2 (left), InSPI2 (middle), and InSPI2 low Mg (right). Shown at the top and right are histograms of each quantity. B) Venn diagram of all unique and shared ON promoters in NonSPI2 (grey, top left), InSPI2 (pink, top right), and InSPI2 low Mg (red, bottom). C) Promoter dynamics in NonSPI2 for the 895 promoters that turned ON in all three media in (B). Each promoter profile (row) was normalized to its own maximum. Color represents the percentage of maximum activity. Profiles were clustered using a centroid-based method to highlight distinct qualitative behaviors (i.e., constitutive, late, pulse). D) Promoter dynamics for the 895 promoters that turned ON in all three media in (B) in NonSPI2 (left), InSPI2 (middle), and InSPI2 low Mg (right). Each promoter profile (row) was normalized to its own maximum. Color represents the percentage of maximum activity. All heatmaps are ordered by ascending *t*_max_ in NonSPI2 to illustrate changes in promoter dynamics in SPI2-inducing media. E) Time to initial activity (*t*_ON_) for virulence-related genes (rows) in InSPI2 (pink), InSPI2 low Mg (red), and NonSPI2 (dark grey). Only promoters that showed activity in all three conditions were included. Reporters were ordered by ascending *t*_ON_ in InSPI2. F) Promoter activity dynamics for six virulence reporters (*ompC*, *sseJ*, *sifB*, *phoP*, *pagK*, *phoN*) in NonSPI2 (grey), InSPI2 (pink), and InSPI2 low Mg (red). Activity is reported as GFP (parent-subtracted to account for autofluorescence) normalized to blanked OD_600_ (to account for changes in cell number). A promoter was defined as ON based on comparison to a dynamic estimate of background noise (**Methods**). ON activity is denoted by solid lines, whereas OFF activity is denoted by dashed lines.

We sought to elucidate the basis of the pulse. Pulsatile behavior still occurred when an antibiotic was added to maintain plasmid selection and in 96-well (rather than 384-well) plate formats (data not shown), suggesting that the decrease in GFP at later times was not due to plasmid loss or poor oxygenation, respectively. We noted that maximum promoter activity in InSPI2 and InSPI2 low Mg occurred approximately coincident with cells entering stationary phase, and we hypothesized that cells underwent a physiological shift during this transition. Single-cell imaging after 24 h of growth in InSPI2 or InSPI2 low Mg revealed cells with disrupted morphologies, whereas cell morphology was normal after 24 h of growth in NonSPI2 (**Fig. S4A**). In InSPI2, after 24 h cells contained regions that were less phase dark, which were likely regions where the outer membrane was separated from the cell wall^35^ (**Fig. S4A**). During growth in InSPI2 low Mg in 12-well plates, cell cultures exhibited lysis after 24 h of growth (**Fig. S4B**), likely due to the role of Mg^2+^ in outer membrane stabilization^36^. Although buffered, pH decreased in all three SPI2 media after 24 h of growth (**Fig. S4C**), which could compound stationary-phase stress. These results reveal that *S.* Tm physiology is strongly impacted by growth in InSPI2, particularly during the transition to stationary phase, and these factors should be considered when evaluating any phenotypes in these media.

This global shift in promoter activity profiles in SPI2-inducing conditions led us to hypothesize that in addition to the known up-regulation of SPI2-related gene activity, there is also time-dependent regulation in response to environmental stimuli, especially prior to cells entering stationary phase. Indeed, for the vast majority of reporters for SPI2-related genes that were active in all 3 SPI2 media, including *ssrA*, *ompC*, *ompF*, *ssaG*, *ssaR*, *sseJ*, *pipB*, *sopD2*, *pipB2*, and *sspH2*, *t*_ON_ was earlier in InSPI2 and InSPI2 low Mg compared with NonSPI2 (**Fig. 2E,F**). Transcription of *phoP*-activated genes is known to be regulated by magnesium levels^3997^, and indeed *t*_ON_ for the *phoP*, *phoN*, *pagC*, and *pagD* reporters was shifted earlier in InSPI2 low Mg compared to InSPI2 (**Fig. 2E,F**). Some other SPI2-related reporters, such as *sifB*, were induced in SPI2-inducing media but not in NonSPI2 (**Fig. 2F**). Taken together, these *in vitro* data demonstrate that our reporter library can capture systems-level changes in activity and dynamic regulation in conditions designed to mimic aspects of the intra-macrophage environment.

### *Salmonella* promoters turn on in multiple stages during macrophage infection

Our *in vitro* data demonstrate that we can capture time-dependent transcriptional regulation, and we next sought to profile such regulation in *S.* Tm within macrophages. To achieve this goal, we infected a macrophage-like cell line with each strain in our library individually and measured intracellular fluorescence via time-lapse fluorescence microscopy coupled to automated cell segmentation. The parent strain (SL1344::*phoN-dTomato*) expresses dTomato within a macrophage, enabling normalization of the GFP signal to bacterial cell density during infection (**Methods**). Intracellular reporter activity was measured using a high-throughput fluorescence microscope (**Methods**). Images were collected from each well in a 96-well black-walled plate in phase and two fluorescence channels (GFP and dTomato) every ∼1 h over 24 hours post infection (h.p.i.). Following automated macrophage segmentation, intracellular reporter activity was quantified for *S*. Tm-positive macrophages, which generated an infection dataset containing >730,000 data points. To classify strains based on phases of the host-pathogen response, we binned infection into four stages: early (0–4 h.p.i.), middle (5–9 h.p.i.), late (10–12 h.p.i.), and escape (13–15 h.p.i.) (**Fig. 3A**).

**Figure 3:**
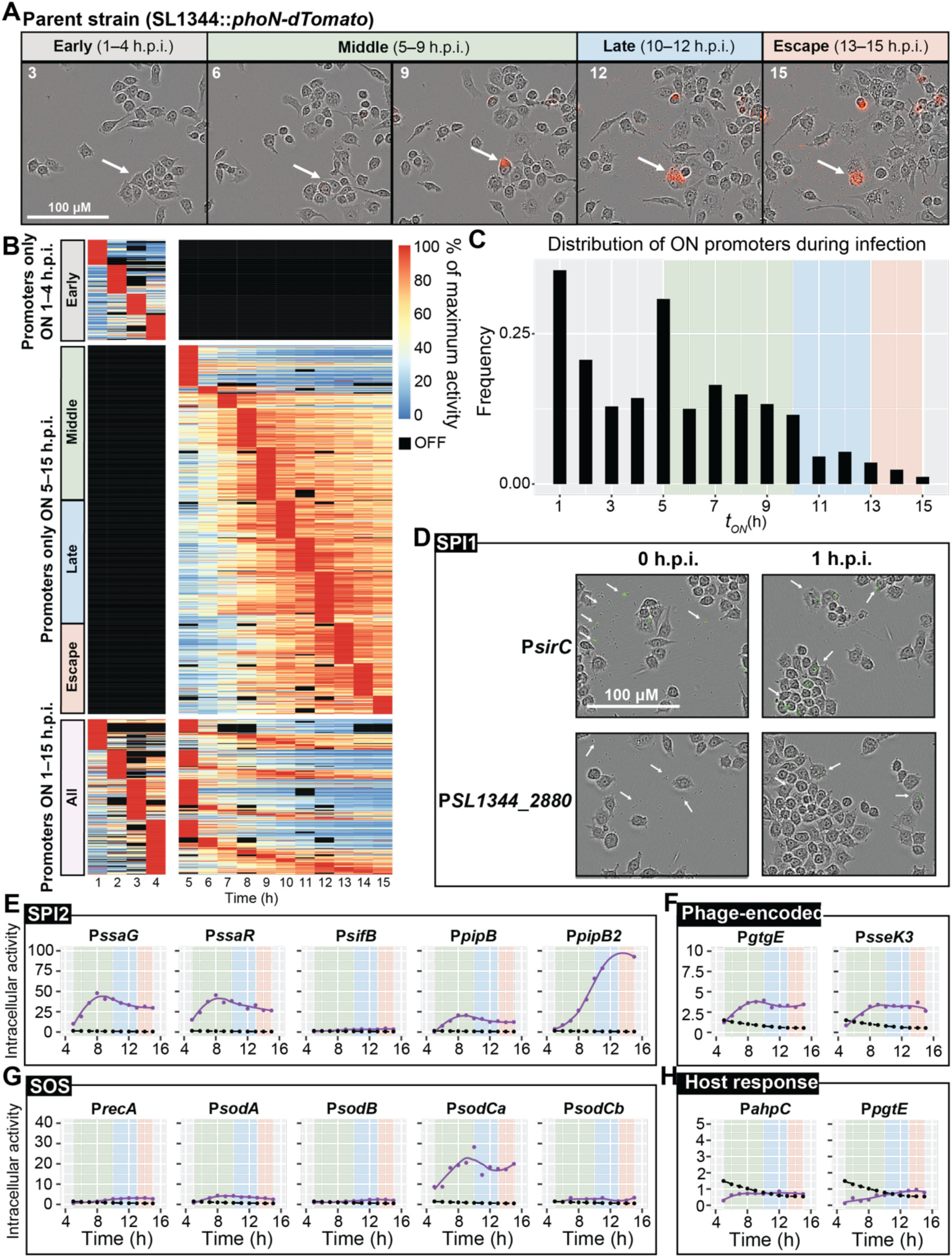
Intracellular *S.* Tm exhibits time-dependent regulation throughout macrophage infection. A) Time-lapse images of the parent strain of the *S*. Tm reporter library, SL1344::*phoN-dTomato*. Shown are phase-contract images overlaid with the dTomato signal, which is produced once *phoN* is induced after responding to SCV maturation. Numbers in the upper left indicate the time in hours. The infection was binned into multiple stages of infection: early (0–4 h.p.i.), middle (5–9 h.p.i.), late (10–12 h.p.i.), and escape (13-–5 h.p.i.). White arrows indicate the same infected macrophage tracked over the 15 h interval. B) Dynamics for 1,007 active promoters in macrophages ordered by ascending *t*_max_. Each promoter profile (row) was normalized to its own maximum. Color represents percentage of maximum activity. Promoters are ordered by ascending *t*_max_ to show that maximum activity is collectively attained across all stages of infection. C) The distribution of times at which activity was initially detected (*t*_ON_) for the 1,007 ON promoters. The background colors for the plots represent the stage of macrophage infection as early (grey), middle (green), late (blue), and escape (orange). D) Time-lapse images of reporters for two promoters in the SPI locus (P*sirC* and P*SL1344_2880*) that are ON during the early stage of macrophage infection. Shown are phase-contrast images overlaid with the GFP and dTomato channels (no dTomato signal was detectable at this early timepoint due to the dependence of *phoN* expression on SCV maturation). Several GFP-positive bacteria were present at the initial time points for pre– and post-invasion (0 and 1 h.p.i., respectively). E) Promoter activity dynamics for several promoters regulating the expression of SPI2-relevant genes between 5–15 h.p.i. Intracellular macrophage activity (parent-subtracted GFP normalized by background-subtracted dTomato) is shown in purple and the dynamic background threshold is shown in black. Points are measurements, and lines are LOESS curve fits. The background colors for the plots represent the stage of macrophage infection as middle (green), late (blue), and escape (orange). F) Activity dynamics for promoters regulating the expression of prophage-encoded genes between 5–15 h.p.i. G) Activity dynamics for promoters that are known to regulate the SOS response between 5– 15 h.p.i. H) Activity dynamics for promoters that reach a maximum in the later stages of infection when *S*. Tm is preparing for escape during host lysis between 5–15 h.p.i.

During infection, a promoter was defined as “ON” based on comparison to a dynamic estimate of background noise over the 1–15 h.p.i. interval (**Fig. S5**). Normalization by the fluorescence signal from the dTomato transcriptional reporter was not performed until 4 h.p.i. since SCV maturation is needed to induce *phoN* expression. Hence, during the early stage of infection (1–4 h.p.i.), we quantified activity of each promoter as the background-subtracted GFP signal (difference between the mean GFP fluorescence for GFP-positive macrophages infected by the strain of interest and the mean GFP fluorescence for all macrophages of the parent strain without a plasmid). Promoter activity during the middle, late, and escape stages (5–15 h.p.i.) was quantified as the background-subtracted GFP signal (difference between the mean GFP fluorescence for dTomato-positive macrophages infected by the strain of interest and the mean GFP fluorescence for all macrophages of the parent strain without a plasmid), normalized by background-subtracted dTomato fluorescence from dTomato-positive macrophages to account for intracellular bacterial replication (**Methods**).

In total, 1,007 promoters were detected as ON during the first 15 h of macrophage infection. There was a wide range of dTomato-normalized GFP signal, and 41% of promoters turned ON prior to 5 h.p.i (**Fig. 3B**). The distribution of time to reach maximum activity (*t*_max_) was multimodal, spanned the 15 h of intracellular infection, and was not correlated with maximum promoter activity (**Fig. S6A**). The distribution of maximum promoter activity was log normal, indicating that most promoters exhibited relatively low activity while a few displayed very high activity (**Fig. S6B**). As expected, the promoter regulating *phoE* (SL1344_0316/SL1344_RS01640), which encodes a phosphate-limitation-inducible outer membrane porin, exhibited the highest activity upon invasion, reflecting the essentiality of the *S*. Tm response to the phosphate-limited intracellular environment^38^ (**Fig. S6C**). We observed promoters turn ON at every hour of infection, reflecting the dynamic programming of *S*. Tm’s response throughout intracellular infection (**Fig. 3C**). The first peak in *t*_ON_ reflects the expected large pulse in promoter activity by invading *S.* Tm to the intracellular environment during the early stage of infection (1–4 h.p.i.)^11^. The distribution of *t*_ON_ values exhibited a second peak at 5 h.p.i. following maturation of the SCV and accumulation of host derived reactive oxygen species (ROS)^8,39,40^ (**Fig. 3C**).

Previous studies documented a precise transcriptional timing pattern for the genes contained within SPIs during macrophage infection^11,13,41^. Specifically, *S*. Tm invasion of macrophages initiates through the upregulation of SPI1 genes that encode the T3SS-1, associated virulence effectors, and regulators^42^. This early stage of infection captures initial *S*. Tm invasion and formation of the SCV^8,43^. Interestingly, prior to invasion several promoters of genes within the SPI1 locus (e.g., *sirC*, *SL1344_2880/SL1344_RS15015*) exhibited high activity when localized near macrophages during the early stage of infection^44,45^ (**Fig. 3D**). We observed that SPI1 promoters exhibited heterogeneity in GFP signal in extracellular *S*. Tm cells (**Fig. 3D**), consistent with previous findings^46,47^. These results show that our library can expand our understanding of heterogeneity across virulence-related promoters.

Following entry of *S*. Tm into macrophages, previous studies showed that acidification of the SCV results in SPI2 upregulation and production of the T3SS-2, which leads to injection of additional effector proteins^48,49^. This middle stage (5–9 h.p.i.) captures the *S*. Tm response to host cell oxidative and nitrosative bursts, maturation of the SCV, and expression of the pathogenicity islands relevant to intracellular replication^39,40^. As expected, the promoters of genes associated with the SPI2 T3SS-2 (e.g., *ssaG*, *ssaR*, *sifB*, *pipB*, *pipB2*) and other virulence-related genes (e.g., *phoP*, *ompC*, *ompF*, *sopD2*, *sseJ*, *sspH2*, *steA*) exhibited high activity during intracellular macrophage infection (**Fig. 3E, S7**). Thus, SPI1 and SPI2 are largely activated sequentially during macrophage infection.

Another common strategy for the genetic acquisition of virulence effectors in *S*. Tm involves phage-mediated integration into the core genome using the content of prophage islands to express effectors required for intracellular replication or survival^50^. However, the precise timing of the transcriptional activation of these loci remains unclear. We observed that the promoters for virulence effectors GtgE (encoded on the Gifsy-2 prophage) and SseK3 (encoded on the ST64B prophage) were active during the middle stage of infection (*t*_ON_=6 h for both)^50,51^ (**Fig. 3F**). Furthermore, the expected timing of ROS accumulation in infected macrophages 5–7 h.p.i.^40^ was followed by increased activity of promoters regulating expression of the SOS response: regulator *recA* (*t*_ON_=8 h), several superoxide dismutases that are important antioxidants for minimizing ROS-mediated damage (e.g., *sodA, t*_ON_= 6 h; *sodB*, *t*_ON_=7 h; *sodCa*, *t*_ON_=5 h; *sodCb*, *t*_ON_=7 h), and other genes related to resistance to oxidative damage (e.g., *ahpF*, *t*_ON_=5 h)^52^ (**Fig. 3G, S8A**).

During the late stage of infection (10–12 h.p.i.), we expected activity from promoters of genes in response to the continued accumulation of ROS in macrophages and ones needed to prime *S*. Tm for the upcoming host lysis and escape^53^. Indeed, we observed genes related to oxidative damage turn ON in the late stage, such as alkyl hydroperoxide reductase^52^ (*ahpC*, *t*_ON_=11 h) (**Fig. 3H**). We also observed activity onset for sigma factor *rpoE* (*t*_ON_=10 h) whose deletion has been found to sensitize *S*. Tm to killing by antimicrobial peptides and ROS^54^ (**Fig. S8A**). The promoter of *pgtE*, which encodes an outer membrane protease responsible for complement cleavage^55^, turned ON at 12 h, likely to prime *S*. Tm for the extracellular environment, (**Fig. 3H**). Additionally, several prophage gene promoters (e.g., *gpE*, *gpO*, *pspA*, and *pspB*) exhibited high activity during the later stages of macrophage infection (**Fig. S8B**).

Lastly, we examined *S.* Tm promoter activity during the escape stage of macrophage infection between 13–15 h.p.i., an interval during which macrophages undergo inflammasome-mediated pyroptosis^56^. The promoter for *yihO*, which encodes a membrane transport protein important for capsule assembly and environmental persistence^57^, turned ON during the later stages of infection (*t*_ON_=13 h), as did *araC,* part of the L-arabinose utilization operon that is important for the carbohydrate metabolism in the gut environment^58^. In summary, mapping the dynamics of transcriptional regulation during macrophage infection corroborated previous findings regarding the early response of *S*. Tm to the intracellular environment^11,59,60^, and provided the precise timing of activation for promoters regulating the expression of virulence-related genes over each stage of infection.

### Comparative analysis between *in vitro* growth and intracellular infection identifies novel infection-relevant promoters

In total, 70.1% (2,016 of 2,874) of the reporter strains were defined as ON in at least one *in vitro* condition and/or during intracellular macrophage infection. By constructing strains for all computationally predicted promoters, our reporter library was able to provide the first experimental validation of some promoters. To generate this list, we first compared to previous RNA-seq results representing a compendium of conditions for potential gene activation, and we manually removed promoters from this list that our literature search identified as confirmed by other methods such as 5’RACE (**Table S4**). Our screens report novel activity for 234 promoters that can be divided into two groups (**Fig. 4A**). In the first group, 112 (out of 283 putative promoters) are considered “newly identified promoters” because they were not included in previous RNA-seq analyses and thus do not have currently available transcriptomics information due to the use of an older (less well annotated) reference genome^12,13^. In the second group, 122 (out of 249) are considered promoters of “newly identified activity” since these regions did not exhibit activity across 22 *in vitro* conditions or during intracellular macrophage infection in previous RNA-seq studies^12,13^. The 234 promoters with novel activity that emerged from our screen encompass a wide range of condition-specific activity across the four *in vitro* media and intracellular infection conditions (**Fig. 4B**) and represent diverse biological processes (**Fig. 4C, Methods**).

**Figure 4:**
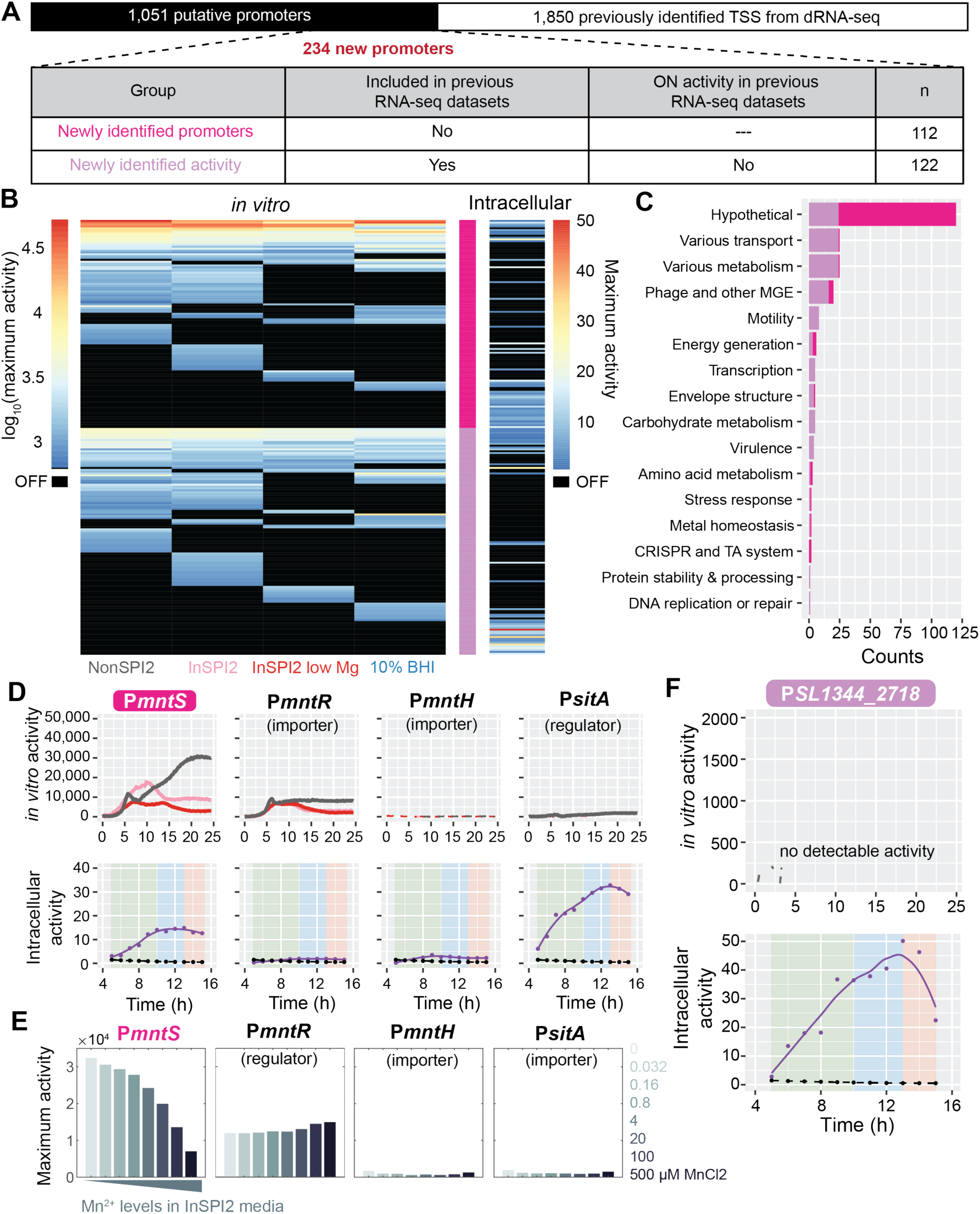
Comparison between *in vitro* growth and intracellular macrophage infection reveals promoters important for manganese regulation. A) Division of the 234 novel promoters identified in our screens into two categories: “newly identified promoters” were not included in previous RNA-seq datasets, and thus do not have currently available expression data (pink); promoters with “newly identified activity” did not show activity in previous RNA-seq datasets across a compendium of *in vitro* and macrophage infection conditions (purple). B) Normalized maximum activity for the 234 putative promoters that were computationally predicted but had not been previously experimentally validated. Maximum activity is shown for the *in vitro* conditions NonSPI2, InSPI2, InSPI2 low Mg, and 10% BHI (left) and intracellular macrophage infection (right). Black values denote OFF. Promoters are grouped by the categories in (A). C) Functional categorization of the 234 novel promoters. D) Promoter activity dynamics for P*mntS*, P*mntR*, P*mntH*, and P*sitA*. P*mntS* is in the “newly identified promoters” group. Top: *in vitro* activity (parent-subtracted GFP normalized by background-subtracted OD_600_) for NonSPI2 (grey), InSPI2 (pink), and InSPI2 low Mg (red). Solid lines denote ON activity, and dotted lines denote OFF activity. Bottom: intracellular macrophage activity (parent-subtracted GFP normalized by background-subtracted dTomato) is shown in purple, and the dynamic background threshold is shown in black. Points are measurements, and lines are LOESS curve fits. The background colors for the plots represent the stage of macrophage infection as middle (green), late (blue), and escape (orange). E) Maximum promoter activity for P*mntS*, P*mntR*, P*mntH*, and P*sitA* in InSPI2 medium supplemented with 0 to 500 µM MnCl_2_. F) Promoter activity dynamics for P*SL1344_2718*, which is in the group of promoters with “newly identified activity.” Top: *in vitro* activity (parent-subtracted GFP normalized by background-subtracted OD_600_) for NonSPI2 (grey), InSPI2 (pink), and InSPI2 low Mg (red). No activity was detected in any *in vitro* conditions. Bottom: intracellular macrophage activity (parent-subtracted GFP normalized by background-subtracted dTomato) is shown in purple, and the dynamic background threshold is shown in black. Points are measurements, and lines are LOESS curve fits. The background colors for the plots represent the stage of macrophage infection as middle (green), late (blue), and escape (orange).

Among the “newly identified promoters,” the reporter for *mntS* (SL1344_RS25090) exhibited high activity in InSPI2 media (high and low Mg) and during intracellular macrophage infection. The promoter region for *mntS* encodes a small (42 aa)^61^ manganese (Mn^2+^) response protein that is predicted to be involved in metal homeostasis, but has not been studied previously in *S*. Tm. In *E. coli*, MntS is important for maintaining Mn^2+^ levels in Mn^2+^-limited environments^62^ by inhibiting the Mn^2+^-exporter MntP^62^. Consistent with previous RNA-seq data^13^, reporters for the Mn^2+^ importer (P*mntH*), Mn^2+^/iron importer (P*sitABCD*), and Mn^2+^ regulator (P*mntR*) exhibited activity during intracellular macrophage infection (**Fig. 4D**). Our findings are consistent with adaptive regulation of manganese levels by *S.* Tm in response to host Mn^2+^ sequestration in the SCV^63^.

Given the *in vitro* and intra-macrophage activity of P*mntS* and other promoters related to Mn^2+^ homeostasis, we hypothesized that P*mntS* would be sensitive to environmental Mn^2+^ levels. Indeed, we found that P*mntS* exhibited a dose-dependent, anti-correlated response to supplemented MnCl_2_ concentration in InSPI2 in a range from 0–500 µM MnCl_2_ (which includes toxic concentrations that are known to activate export of Mn^2+^ through MntP and YiiP^64^) (**Fig. 4E**). These findings are consistent with a previous study showing that MntS protein levels and promoter expression levels are anti-correlated with environmental Mn^2+^ concentrations in *E. coli*^61,65^. Our *mntP* and *yiiP* reporters did not show signal in any experiment (data not shown), likely because these reporters are non-functional. Nonetheless, because P*mntS* activity was more sensitive to Mn^2+^ levels than the regulator P*mntR* or importers P*mntH* and P*sitA*, we speculate that MntS plays an important role in sensing and regulating intracellular Mn^2+^ levels in *S*. Tm.

Furthermore, our data set provided the first reported activity of many other promoters, 96 of which are annotated with hypothetical functions (**Fig. 4C**, **Table S4**). Among these promoters with “newly identified activity,” we observed relatively high activity of the SopE-Phi prophage-associated *P*SL1344_2718 during intracellular infection (**Fig. 4F**). Phyre2^66^ predicted the structure of SL1344_2718 to be a phage capsid protein with 100% confidence and 96% coverage. Interestingly, this promoter did not turn ON in any of the *in vitro* conditions, illustrating a case of macrophage-specific induction (**Fig. 4F**). These findings establish that our library is a powerful tool for identifying promoters with previously unrecognized infection relevance, adding to the repertoire of genes necessary for *Salmonella* to survive and replicate in the SCV environment.

### Systems-level functional analysis reveals a metabolic pathway important for intracellular macrophage infection

Our identification of new promoter activity during infection, coupled with time-dependent intra-macrophage transcriptional regulation, suggested the potential to discover novel regulatory patterns crucial for intracellular survival. To determine links between promoter activity dynamics and *S*. Tm pathogenesis, we conducted a functional characterization of the 1,007 promoters induced during macrophage infection. The two largest groups involved promoters that regulate the expression of hypothetical and metabolism-related genes (**Fig. S9)**. Other pathways enriched during macrophage infection include transport, virulence, and the redox response, consistent with previous findings^13,41,60^. This analysis also highlighted the carbohydrate metabolism functional group, which includes promoters of genes in the Entner-Doudoroff (ED) metabolic pathway including *kdgK*, *kdgT*, *edd*, *eda*, *idnK*, *uxuA*, and *gnd*. The normalized fluorescence signal of several of these promoters first displayed activity between 8–15 h.p.i. (**Fig. 5A**). Since it is known that the ED pathway is upregulated during macrophage infection^11,21^ and utilized to increase metabolic flux^67^, we investigated the time-dependent regulation of this pathway. GFP fluorescence from the P*kdgK* reporter strain during macrophage infection increased after the increase in P*phoN*-regulated dTomato expression that reflected SCV maturation^8^ (**Fig. 5B**).

**Figure 5:**
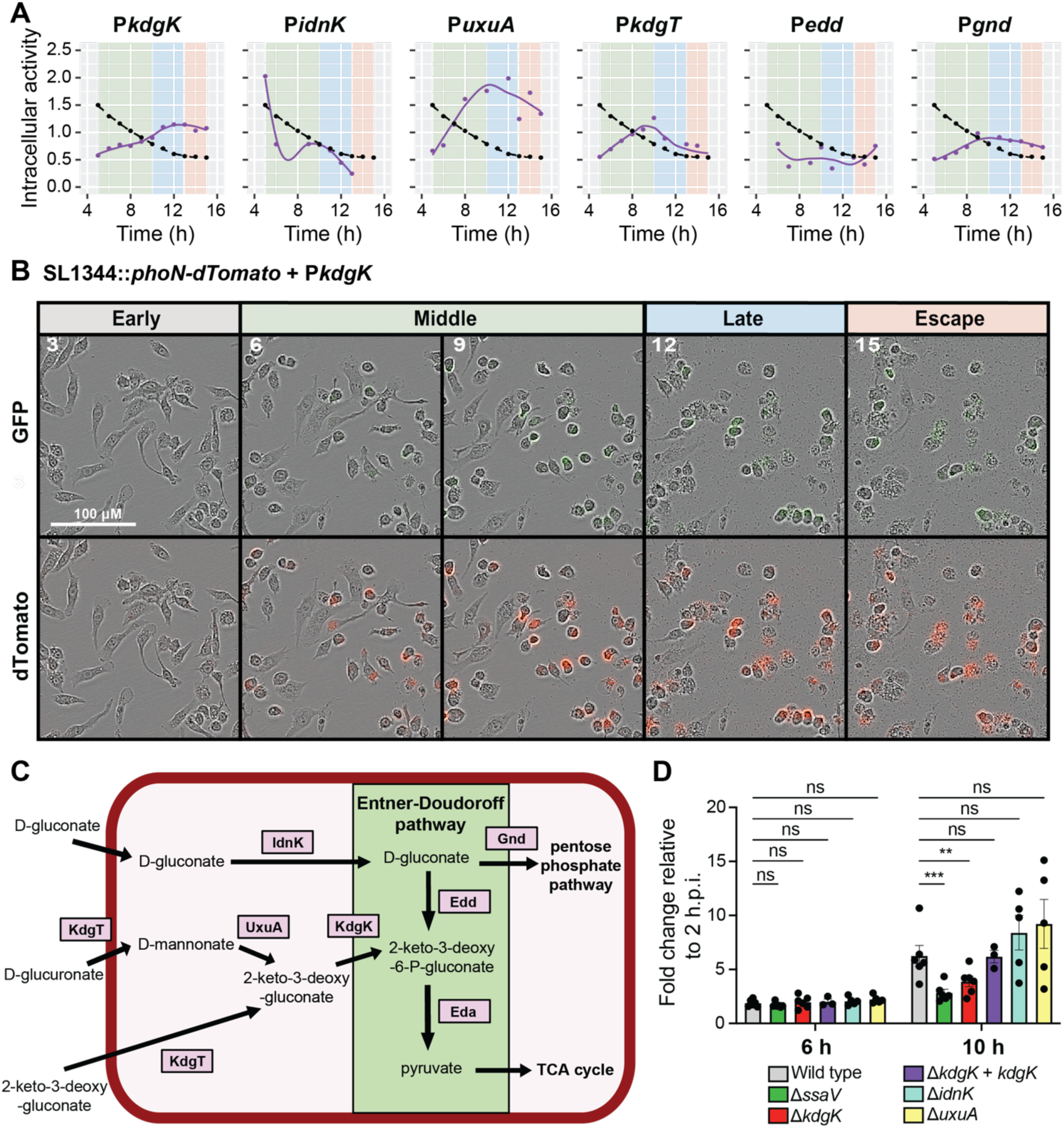
*S*. Tm depends on the Entner-Doudoroff (ED) pathway during later stages of macrophage infection. A) Normalized GFP dynamics from *S*. Tm in infected macrophages for several promoters regulating the expression of genes related to the ED pathway (*kdgT*, *kdgK*, *edd*, *uxuA*, *idnK*, *gnd*) between 5–15 h.p.i. Intracellular activity is parent-subtracted GFP normalized by background-subtracted dTomato. Points are measurements and lines are LOESS 5 B) ure curve fits. Each reporter was compared to the background threshold (black dotted line). The background colors for the plots represent the stage of macrophage infection: middle (green), late (blue), and escape (orange). C) Time-lapse images of the activity of the P*kdgK* reporter strain during macrophage infection. Shown are phase-contrast images overlaid with the GFP (top) or dTomato (bottom) signal. Numbers in the upper left indicate the time in hours. D) Catabolic pathways of several sugar acids including the ED pathway (highlighted in green). E) Macrophages were infected with *S*. Tm mutant strains carrying the plasmid pFCcGi. Macrophages were harvested, fixed, and stained at 2, 6, and 10 h p.i. for analysis by flow cytometry. Fold replication was determined by comparing the ratio of mean GFP to mean mCherry signal at 6 and 10 h.p.i. to the ratio at 2 h.p.i. for each mutant. Error bars represent 1 standard error of the mean (SEM) for *n*=5 independent experiments for each mutant and *n*=3 independent experiments for the Δ*kdgK*+*kdgK* complementation strain. Multiple unpaired t-tests were conducted for each strain relative to wild type with a two-stage step-up method for multiple hypothesis correction. **: *p*<0.01; ***: *p*<0.001.

Since the ED gene promoters exhibited increased activity in the middle, late, and escape stages of infection, we hypothesized that the ED pathway is important for intracellular *S*. Tm survival. The ED pathway is comprised of the enzymes 6-phosphogluconate dehydratase (encoded by *edd*) and KDGP aldolase (encoded by *eda*). The pathway culminates in aldol cleavage of 2-keto-3-deoxy-6-phosphogluconate (KDPG) into pyruvate to funnel into the tricarboxylic acid (TCA) cycle, ultimately contributing to energy production^68^ (**Fig. 5C**). Consistent with our findings, a previous study using a *S.* Tm GFP reporter observed increased expression from the *edd* promoter within RAW macrophages^21^. Furthermore, isotope tracing of *S*. Tm metabolism during macrophage infection revealed high metabolic flux through the ED pathway, which converts 6-carbon sugars (e.g., hexuronates, gluconate, glucuronate, and galacturonate) into KDPG^67^.

To validate the importance of the ED pathway during intracellular *S*. Tm replication, we individually deleted the *kdgK*, *idnK*, and *uxuA* genes, which encode enzymes that feed metabolites into the ED pathway. We measured bacterial replication rate using a previously developed dual-reporter plasmid (pFCcGi)^69^. *S*. Tm Δ*kdgK*, Δ*idnK*, and Δ*uxuA* strains exhibited replication and growth rates comparable to wild type across *in vitro* conditions, as did a Δ*kdgK* complementation strain. The replication rate in macrophages was measured at 2, 6, and 10 h.p.i. using flow cytometry. The mean fold-change in replication rate at 6 h.p.i. relative to 2 h.p.i. was similar (∼1.6–1.8x increase) across all strains (**Fig. 5D**), while a Δ*ssaV* strain deficient in SPI2 T3SS replicated significantly less than wild type in macrophages, as previously reported^70^. The Δ*idnK* and Δ*uxuA* strains, which lack genes encoding a D-gluconate kinase and mannonate dehydratase, respectively, exhibited high variance in intracellular replication but were not significantly different from wild type. Importantly, deletion of *kdgK*, which encodes an enzyme that catalyzes KDPG production, resulted in significantly lower levels of replication at 10 h.p.i. compared to wild type; this defect was rescued by complementation of *kdgK* at the native locus (**Fig. 5D**). These results suggest that the ED pathway is important for the survival and replication of *S*. Tm by compensating for nutrient limitation in the macrophage environment at later stages of infection. More generally, these findings demonstrate the importance of profiling the dynamics of transcriptional activity, particularly within changing conditions such as the intracellular environment during infection.

## Discussion

Fluorescence reporters are powerful tools for quantification of transcriptional dynamics at high temporal resolution, an especially relevant goal for bacteria like *S*. Tm that rapidly adapt their programming during environmental shifts. Capturing the dynamics across all stages of intracellular macrophage infection revealed a metabolic shift during the later stages of infection involving the ED pathway (**Fig. 5D**). Future investigations into the metabolic profile of intracellular *S*. Tm should involve integrating metabolic information from the host as well as the pathogen^67^, including consideration of the time dependence of nutrient utilization. Mammalian cells employ multiple primary types of cellular metabolism. For example, when macrophages are activated in response to bacterial pathogens, they can undergo a shift from mitochondrial oxidative phosphorylation to aerobic glycolysis leading to depletion of intracellular glucose stores and the availability of oxygen, a typical terminal electron acceptor for aerobic metabolism in bacteria^15^. Importantly, these processes have been shown to be partly dependent on macrophage polarization status and need further exploration in this setting^71^. We propose that the host undergoes changes in metabolism that limit intracellular *S*. Tm replication through nutrient sequestration. In turn, *S*. Tm shifts its own metabolic programming at later stages of infection to overcome the glucose-limiting conditions in the SCV. This model is consistent with our findings wherein the ED pathway may provide alternate carbon sources to support *S.* Tm replication, although the replication defect of the Δ*kdgK* strain could also result from accumulation of toxic metabolic intermediates such as KDGP^72^. Taken altogether, our work provides further support for dynamic metabolic crosstalk between host and pathogen and motivates further exploration of the potential direct or indirect effects of ED pathway metabolic by-products on the host cell environment and immune responses.

Our library screens provided the first evidence of activity for many *S.* Tm genes, including some involved in pathogenicity (**Fig. 4B-F**). Previously published RNA-seq datasets catalogued conditions under which many *S*. Tm genes are induced and identified TSS using dRNA-seq^12,13^. Our library and analyses complement these data sets by identifying activity from novel promoter regions. Some of the novel promoter identifications result from our use of a genomic reference with updated annotations and an unbiased computational approach to identify promoter regions. Our ability to identify activity from some chromosomal regions whose expression was not detected across 23 growth conditions using RNA-seq is likely to reflect our temporal approach that spanned both growth and infection stages. Overall, our findings provide insight into the regulatory regions of the *S*. Tm genome and highlight potential targets at different stages of infection for future drug discovery focused on the inhibition of *S.* Tm pathogenesis.

Our comparative analysis among expression profiles during intracellular infection and controlled *in vitro* growth highlights the power of our library to uncover the infection relevance of novel promoter regions. In particular, our data sets revealed increased activity of several promoters (e.g., P*mntR*, P*zur*, P*fur*) that regulate response of *S*. Tm to host nutritional immunity through metal ion sequestration (**Fig. S10**). Our screens led to the hypothesis that *mntS* regulates the *S.* Tm response to metal-limiting conditions in the SCV, and we found that P*mntS* activity was dependent on environmental Mn^2+^ concentration *in vitro* (**Fig. 4E**). Because the activity of known Mn^2+^ regulators, importers, and exporters lacked the sensitivity to Mn^2+^ concentration of P*mntS*, we hypothesize that MntS plays a critical role in the ability of *S*. Tm to sense Mn^2+^ limitation in the SCV (**Fig. 4E**). Moreover, our study highlights the remaining knowledge gap about *S.* Tm adaptation during intracellular infection. Our candidate list of 234 novel promoters, many with hypothetical or unknown functional annotations, and their quantified expression profiles should provide compelling inspiration and a useful resource for future studies.

Our analyses also revealed that many prophage-associated promoters were activated during infection (**Fig. 4F, S9**). One explanation for this activity is upregulation of genomic loci containing phage-encoded virulence factors^50,51^ (**Fig. 3F**). We also observed activity for other promoters of prophage-associated genes, including ones involved in structural components and excision, during the later stages of infection. This late-stage prophage promoter activity may be the result of accumulation of DNA damage following ROS exposure during intracellular *S*. Tm replication and the resulting SOS response, a known inducer of prophage loci^50,73–75^. While further investigation is needed to elucidate whether host ROS production can lead to prophage induction, these data may provide insight into other mechanisms of *Salmonella* survival during infection. For example, in the intracellular pathogen *Listeria monocytogenes*, its DNA uptake competence (Com) system is required during intracellular infection to promote escape from the macrophage phagosome, and regulation of the Com system relies on prophage excision^76^.

Promoter constructs have specific limitations for transcriptional profiling. For example, our bioinformatic approach may not accurately capture the promoters for every coding region. Additionally, some promoter regions that were expected to have high activity in SPI2-inducing conditions (e.g., P*ssaM*) did not show activity either *in vitro* or during macrophage infection, potentially due to the lack of key regulatory regions that lie >350 bp upstream of the TSS. In our analyses, we focused on the large fraction of reporter strains that showed activity in at least one condition, but false negative reporters could be used to investigate such regulation through further strain construction (e.g., with larger upstream regions^77^). We opted to use a fast-folding and stable GFP to accurately capture the first initiation of activity, hence GFP fluorescence signal does not provide an accurate estimate of downregulation of promoter activity or translational regulation. We note that the absence of complete repressor-binding sites could drive artificial promoter activity. We discovered that SPI2-inducing *in vitro* conditions impose physiological defects during stationary phase, and it is unclear whether such changes reflect conditions in the SCV. This work, combined with other RNA-seq datasets, can be used to improve *in vitro* media conditions designed to mimic aspects of the intra-macrophage environment.

Our bulk measurement approach to profiling transcriptional dynamics leaves outstanding questions about phenotypic heterogeneity. In this work, we demonstrate heterogeneous expression of SPI2 genes such as *ssaR* and *ssaG* (**Fig. 1G,H**). As expected, we also observed qualitative heterogeneity for SPI1 promoters during the early stages of macrophage invasion (**Fig. 3D**), and our library should serve as a powerful tool to expand our knowledge of which genes exhibit heterogenous expression and to understand how *S*. Tm regulates bistable expression during infection^47^ using flow cytometry and high-throughput imaging^78^. By employing higher imaging resolution and improved single-cell segmentation, our library can be used to quantify the activity and heterogeneity of extracellular and intracellular bacteria during invasion of and replication in host cells (**Fig. 3D**)^79^. Moreover, our infection data only includes time points up to host cell lysis due to the increase in noise and inability to perform accurate image segmentation; nonetheless, our library provides future opportunities to profile and discover *S.* Tm gene regulation that occurs during host cell lysis and bacterial escape that could be important for pathogenesis.

Much remains to be discovered about *S.* Tm pathogenesis. The substantial set of genes whose expression has not yet been observed motivates screening across a more extensive and diverse collection of conditions to provide insight into gene functions. Our data can be used to inform time point selection for future RNA-seq or RT-qPCR experiments. Our library could also be used to infect different immune cell types or study the effects of various host-signaling factors to understand host-specific virulence programming. To probe *S.* Tm colonization throughout the gastrointestinal tract, the library could be screened *in vitro* in conditions that include bile, short chain fatty acids (SCFAs), gut-relevant carbon sources, and interactions with other gut commensals. To assist in the identification of therapies against *Salmonella* infections, future screens of the promoter library should include antibiotics, small molecules, and bacteriophages.

Our library of donor *E. coli* strains for high-throughput conjugation also enables straightforward plasmid transfer into other *Salmonella* strains or serovars. Collectively, our existing data and the future applications of our promoter libraries should provide a systems-level understanding of the timing of transcriptional programming during intracellular infection, with the potential to discover virulence mechanisms and treatments with global health impact.

## Acknowledgements

We thank Marlin Jenkins (U. Minnesota) for providing the *S*. Tm SL1344::*phoN-dTomato* strain for library construction, Dirk Bumann (U. of Basel, Biozentrum) for providing the pFOK plasmid for site-specific mutagenesis, Sophie Helaine (Harvard Medical School) for providing the pFCcGi plasmid for quantifying bacterial replication, and Phoom Chairatana (Stanford) for providing guidance on plasmid construction. This study was funded in part by Award T32-AI007328-36 from the National Institute of General Medical Sciences (to O.R.D.), the Blavatnik Family Fellowship Fund (to O.R.D.), an NSF Graduate Research Fellowship (to T.H.N.), NSF Awards EF-2125383 and IOS-2032985 (to K.C.H.), NIH Awards R01 AI147023 and RM1 GM135102 (to K.C.H.), the Allen Discovery Center for Systems Modeling of Infection (to K.C.H., and D.M.M.), grants R01-AI116059 and R01-AI095396 from the National Institute of Allergy and Infectious Diseases (to D.M.M.), and Wellcome Trust Senior Investigator Award 222528/Z/21/Z (to J.C.D.H.). For the purpose of open access, the author has applied a CC BY public copyright license to any Author Accepted Manuscript version arising from this submission. The contents of this study are solely the responsibility of the authors and do not necessarily reflect the views of the Paul G. Allen Institute or the National Institutes of Health. The funders had no role in study design, data collection, and interpretation, or the decision to submit the work for publication.

## Author Contributions

M.R., K.C.H., and D.M.M conceptualized research. T.H.N., O.R.D., D.M.M., and K.C.H. designed the research. M.R. performed computational identification of promoter regions. T.H.N., O.R.D., M.R., D.S.C.B., and B.X.W. performed the research. T.H.N. and O.R.D. analyzed the data.

J.C.D.H. provided supervision. T.H.N., O.R.D., D.M.M., and K.C.H. wrote the paper, and all authors reviewed the manuscript prior to submission.

## Data and code availability

All data are available at https://purl.stanford.edu/fc012fq3845. Custom code used in this paper is available at https://doi.org/10.5281/zenodo.8339637. Any additional information required to reanalyze the data reported in this paper is available from the lead contact upon request.

## Methods

### Bacterial strains, macrophage cell lines, and growth conditions

*Salmonella enterica* serovar Typhimurium (*S*. Tm) strain SL1344 with *phoN*-*dTomato* integration (SL1344::*phoN*-*dTomato*) was used to construct the promoter library and for site-specific mutagenesis. *S.* Tm and *E. coli* strains used for conjugation are listed in **Table S1**. In preparation for screens, bacterial strains were grown overnight in selective media supplemented with 20 µg/mL chloramphenicol (Cm) and/or 50 µg/ml streptomycin (Strep) as required. For all intracellular infection studies, RAW 264.7 murine macrophages (ATCC #TIB-71) were maintained at 37 °C with 5% CO_2_ in Dulbecco’s Minimal Essential Medium (DMEM) (Invitrogen #11995073) supplemented with 10% Fetal Bovine Serum (Fisher Scientific #26-140-079).

### Computational identification of promoter regions

2,907 lead-operon reporter regions were computationally identified based on open reading frames, intergenic regions, and experimentally validated transcriptional start sites^13^ in the genome of *S.* Tm SL1344 (NCBI reference sequence NC_016810.1). Promoter regions were defined as the 350 bp upstream of and including the translational start site (353 bp total). Our library includes intergenic regions longer than 40 bp. The low-copy plasmid backbone pUA66^20^ was used to construct the library with mGFPmut2^18^, a *Cm^R^* cassette, and *mob* genes for conjugative transfer from pSC101. For reporter plasmid constructs, each promoter was fused to *mGFPmut2* (including the S65A, V68L, S72A, and A206K mutations)^18^. Plasmids were assembled, sequence verified, and arrayed into 96-well plates by Thermo Fisher Scientific. All promoter sequences and respective well locations are available in **Table S2**, including promoter sequences that were not successfully cloned.

### High-throughput plasmid transformation into *E. coli*

For each promoter plasmid, 50 µL of competent *E. coli* (MG1655 MFDpir RP4-2-Tc:[*Mu1*::*aac*(3)IV-Δ*aphA*-Δ*nic35* Δ*Mu2*::*zeo*] Δ*dapA*::(*erm*-*pir*) Δ*recA*) cells were mixed with ∼10 ng of plasmid in one well of a 96-well PCR plate (Bio Rad #MLL9601). The mixtures were exposed to a cold shock via incubation on ice for 30 min, then heat shock in a 42 °C water bath for 45 s, followed by another cold shock on ice for 5 min. Cells were then added to 500 µL of fresh Lennox broth (LB, Fisher Scientific #50488761) containing 0.3 mM diammonium phosphate (DAP) in a deep-well 96-well plate (USA Scientific #1896-2110). Plates were sealed with breathable seals (Excel Scientific #LMT-AERAS-EX, T896100-S) and recovered for 1 h at 37 °C with shaking. To select for positive transformants through liquid selection, an additional 500 µL of LB containing 0.3 mM DAP and 40 µg/mL Cm (for a final Cm concentration of 20 µg/mL) were added and cells were grown overnight with shaking at 37 °C. In preparation for high-throughput conjugation, *E. coli* donor transformants were diluted 1:200 into fresh LB containing 0.3 mM DAP in 96-deepwell plates for another round of Cm (20 µg/mL) selection. After overnight growth with shaking at 37 °C, cultures were stored with 25% (v/v) glycerol at –80 °C in 96-well flat-bottom plates (Greiner Bio-One #655161) sealed with aluminum seals (Thermo Fisher Scientific #12-565-398).

### High-throughput conjugation from E. coli to S. Tm

In preparation for high-throughput conjugation, an *E. coli* (donor) culture was grown under selection with 20 µg/mL Cm and a *S.* Tm (recipient) culture was grown under selection with 50 µg/mL Strep (natural resistance) overnight in LB with shaking at 37 °C. The overnight cultures were pelleted, washed, resuspended at 10X density, and mixed at a 1:4 (donor:recipient) ratio. Using a Benchsmart 96 semi-automatic pipetting system (Rainin), 5 µL of conjugation reactions were pipetted onto a rectangular LB-agar plate (Thermo Fisher Scientific #267060) containing 0.3 mM DAP and incubated at 37 °C for 5 h. Colonies were resuspended in LB and serially diluted on selective rectangular LB-agar plates (Thermo Fisher Scientific #264728) containing 20 µg/mL Cm and 50 µg/mL Strep without DAP to screen for positive *S*. Tm transformants and select against *E. coli* donor cells, respectively. After overnight growth at 37 °C, single colonies were picked, grown to saturation at 37 °C in LB containing 20 µg/mL Cm and 50 µg/mL Strep, and stored with 25% glycerol at –80 °C in 96-well flat-bottom plates (Greiner Bio-One #655161).

### Sequence verification of the promoter region in *S.* Tm strains

*S.* Tm strains were struck on LB-agar plates containing 20 µg/mL Cm and 50 µg/mL Strep and grown overnight at 37 °C. Colony PCR was performed by combining 25 µL of Accustart II 2x SuperMix (Quantabio #95137), 1 µL of 10 mM forward primer (PromLib_FWD 5′-AATAGGCGTATCACGAGG-3′), 1 µL of 10 mM reverse primer (PromLib_REV 5′-CCATCTAATTCAACAAGAATTGGG-3′), 18 µL of nuclease-free water, and 5 µL of a single colony diluted into 100 µL of nuclease-free water. The PCR program was 94 °C for 3 min, 35 cycles of [94 °C for 45 s, 50 °C for 60 s, and 72 °C for 90 s], followed by 72 °C for 10 min. Amplified products were confirmed by Sanger sequencing (Elim Bio).

### In vitro screening of the reporter library

For *in vitro* screens, four media were used: NonSPI2 (SPI2 non-inducing), InSPI2 (SPI2 inducing), InSPI2 low Mg, and 10% BHI. Phosphate-carbon-nitrogen (PCN) base medium was used to make NonSPI2 (pH 7.4, 25 mM P_i_, 1 mM MgSO_4_), InSPI2 (pH 5.8, 0.4 mM Pi, 1 mM MgSO_4_), and InSPI2 low magnesium (pH 5.8, 0.4 mM Pi, 10 µM MgSO_4_)^12,28^. 10% Brain Heart Infusion (BHI, BD #2237500) medium was 100% BHI diluted in M9 salts (Sigma-Aldrich #M9956) and supplemented with 0.1 mM CaCl_2_ and 2 mM MgSO_4_; 10% BHI was used instead of 100% BHI to reduce background autofluorescence and thereby improve the dynamic range of GFP measurements. More details about media conditions can be found in **Table S5,6**. *S.* Tm glycerol stocks were pinned on selective rectangular LB plates (Thermo Fisher Scientific #267060) containing 20 µg/mL Cm and 50 µg/mL Strep. After overnight growth at 37 °C, colonies were picked and grown overnight at 37 °C in 200 µL of NonSPI2 or 10% BHI medium containing 20 µg/mL Cm and 50 µg/mL Strep in flat-bottom 96-well plates (Greiner Bio-One #655161) sealed with breathable seals (Excel Scientific #LMT-AERAS-EX, T896100-S). Using a Benchsmart 96 semi-automatic pipetting system (Rainin), overnight cultures were diluted 1:100 in 80 µL of the appropriate medium (NonSPI2 into NonSPI2, InSPI2, or InSPI2 low Mg, or 10% BHI into 10% BHI) in black-walled, clear-bottom 384-well plates (Greiner Bio #781097). Plates were sealed using transparent seals (Excel Scientific #STR-SEAL-PLT) with small, laser-cut holes (∼0.5 mm) for gas exchange. Growth curves were measured using a Biotek Synergy H1 with continuous shaking at 37 °C for 24 h, during which OD_600_ and GFP (488 nm/520 nm excitation/emission) were measured every 10 min.

### In vitro screening *data analysis*

To quantify promoter expression, the GFP signal of the parent strain (no plasmid control) was subtracted to correct for background fluorescence. To normalize for cell number, the parent-subtracted GFP signal was normalized by OD_600_ after subtracting the background absorbance of a blank well (no cells). In all analyses, the denominator was set to a minimum value of 0.3 to avoid fluctuations resulting from division by small values. A promoter was classified as ON if its expression was at least two standard deviations above a dynamic estimate of background noise for at least 3 timepoints (30 min) (**Fig. S2**). Background noise was estimated by calculating the mean over a time interval *t\** of *n\** of the lowest-expressing promoters that could be safely assumed to be OFF in the medium of interest. For each medium, *t\** was identified as the time range starting when OD_600_ reached 0.3 and ending when promoter activity was still detectable in each condition to avoid the physiological complications of late stationary phase in InSPI2 and InSPI2 low Mg. Therefore, *t\** was identified for NonSPI2 as 8–24 h, 8–16 h for InSPI2, 8–16 h for InSPI2 low Mg, and 4–24 h for 10% BHI. To calculate *n\**, the expression dynamics of the *n* lowest-expressing promoters were averaged for a range of values of *n*, using only data within *t\**. Finally, the value of *n\** was selected based on the mean trajectory being closest to 0. All data analyses were performed in R with custom scripts, using packages *dplyr*, *genefilter*, *ggExtra*, *ggplot2*, *gridExtra*, *numbers*, *numDeriv*, *pheatmap*, *Rcolorbrewer*, *realxl*, *svglite*, *tidyverse*, *VennDiagram*, and *zoo*. All data are available at https://purl.stanford.edu/fc012fq3845. All custom scripts are available at https://doi.org/10.5281/zenodo.8339637.

### Single-cell fluorescence imaging and analysis

Single colonies of reporter and parental strains were inoculated into 200 µL of NonSPI2 medium. After overnight growth at 37 °C, saturated cultures were diluted 1:100 into 200 µL of fresh NonSPI2, InSPI2, and InSPI2 low Mg media and were grown with shaking at 37 °C. To aid in image segmentation, all cultures were diluted 1:5 in PBS at 8 h, and 1:20, 1:10, and none for NonSPI2, InSPI2, and InSPI2 low Mg, respectively, at 24 h. For imaging, 1 µL of cells was spotted on a 1.5% PBS-agarose pad and allowed to dry before sealing with a coverslip. Phase-contrast images were acquired with a Ti-E inverted microscope (Nikon Instruments) using a 100X (NA: 1.40) oil immersion objective and a Neo 5.5 sCMOS camera (Andor Technology). Images were acquired using µManager v. 2.0. To calculate GFP intensity per cell, the MATLAB image processing package *Morphometrics*^80^ was used to segment cells from phase-contrast images. Images were filtered for single-cell contours. For each of >1,000 contours in each condition, the median background-subtracted GFP fluorescence was calculated for each cell. Phase-contrast and GFP images were overlaid in FIJI. All data analysis was performed with custom MATLAB scripts. All data are available at https://purl.stanford.edu/fc012fq3845. All custom scripts are available at https://doi.org/10.5281/zenodo.8339637.

### pH measurements during in vitro growth

A single colony of the parent strain was inoculated into 5 mL of NonSPI2 containing 50 µg/mL Strep and grown overnight with shaking at 37 °C. Overnight cultures were diluted 1:100 into 5 mL of fresh NonSPI2, InSPI2, or InSPI2 low Mg in three technical replicates. After 24 h of growth, cultures were centrifuged at 4,000*g* for 10 min and filter sterilized with 0.22-µm filters. The pH of NonSPI2, InSPI2, and InSPI2 low Mg cultures before and after 24 h of *S.* Tm growth was measured using a pH probe (Thermo Fisher Scientific #13-620).

### Intracellular screening in macrophages

To characterize transcriptional responses of *S.* Tm during infection, we performed high-throughput, time-lapse fluorescence microscopy in RAW 264.7 murine macrophages. For infection, macrophages were seeded in black-walled, clear-bottom 96-well plates (Corning #3603) at ∼10^4^ cells per well. *S.* Tm glycerol stocks were pinned onto selective rectangular LB-agar plates (Thermo Fisher Scientific #267060) containing 20 µg/mL Cm and 50 µg/mL Strep. After overnight growth at 37 °C, colonies were picked into 500 µL of LB with 20 µg/mL Cm and 50 µg/mL Strep in round-bottom, 96-well deep-well plates (VWR #76210-518), sealed with breathable seals (USA Scientific #9126-2100), and grown for 16 h at 37 °C. *S.* Tm cultures were washed in PBS (Invitrogen #10010-049) and resuspended in FlouroBrite DMEM (FDMEM, ThermoFisher #A1896701) supplemented with 7% Fetal Bovine Serum, 10 mM HEPES (Gibco #15630-080), 2 mM L-glutamine (Thermo #25030081), and 500 µg/mL L-histidine (Sigma #H6034-25G). Macrophages were infected for 40 min at a multiplicity of infection of 10:1 (bacteria:macrophages) and centrifuged at 250*g* for 5 min. Following the invasion period, cultures were maintained in FDMEM supplemented with 100 µg/mL gentamicin for 50 min to kill extracellular bacteria. Infected macrophages were washed with PBS and maintained in FDMEM supplemented with 15 µg/mL gentamicin for 24 h post infection. Plates of infected macrophages were imaged in triplicate per well in an Incucyte S3 Live-Cell Analysis Platform (Sartorius #4647). Phase and fluorescence images were collected every hour per well using a 20X objective. Fluorescence images were acquired using red (excitation: 565-605 nm, emission: 625-705 nm) and green (excitation: 440-480 nm, emission: 504-544 nm) channels.

### Macrophage infection image analysis

Fluorescence background subtraction was performed using the IncuCyte Surface Fit algorithm and cell outlines were segmented using the Cell-by-cell analysis software (Sartorius #9600-0031). After segmentation, cells were classified into GFP OFF/ON and dTomato OFF/ON according to fluorescence intensity thresholds (GFP: 0.0853 arbitrary fluorescence units (AFU), dTomato: 0.0309 AFU). At each time point between 1–4 h.p.i., promoter activity was quantified as the difference between the mean GFP fluorescence for GFP-positive macrophages infected by the strain of interest and the mean GFP fluorescence for all macrophages of the parent strain without a plasmid to correct for background and fluorescence bleed-through. At each time point between 5–15 h.p.i., promoter activity was quantified as the difference between the mean GFP fluorescence for dTomato-positive macrophages infected by the strain of interest and the mean GFP fluorescence for all macrophages of the parent strain without a plasmid to correct for background and fluorescence bleed-through. To normalize for bacterial cell number, the background-subtracted GFP signal was normalized by the dTomato signal of the promoter strain after subtraction of the background dTomato signal of a separate well seeded with uninfected macrophages. In all analyses, minimum values of 0.005 AFU and 0.04 AFU were used for the GFP and dTomato signals, respectively, to avoid fluctuations resulting from division by small values. Following normalization, fluorescence signal is reported as relative fluorescence units (RFU). A dynamic estimate of background noise for each time window was determined as the mean GFP signal from GFP-(1–4 h.p.i.) or dTomato-positive (5–15 h.p.i.) macrophages infected with the parent strain for each time point. A promoter was classified as active if the GFP signal was at least two standard deviations above a dynamic estimate of background noise for a given time window (**Fig. S6**). All data analyses were performed in R using custom scripts. All data are available at https://purl.stanford.edu/fc012fq3845. All custom scripts are available at https://doi.org/10.5281/zenodo.8339637.

## Functional annotations

Functional annotations were obtained for the *S.* Typhimurium LT2 reference genome, leveraging existing Gene Ontology (GO) terms for biological processes and molecular functions that were exported from BioCyc. These annotations were subsequently mapped to the *S.* Typhimurium SL1344 genome through ortholog mapping. To streamline genome-wide functional annotations, BioCyc-based annotations were subjected to manual curation, resulting in a total of 91 categories (**Table S2**). Unnamed genes with no recorded GO terms were designated as hypothetical genes. Subsequently, the functional category for each promoter designated as ON in the dataset was recorded to quantify the number of promoters belonging to each functional class.

### *S.* Tm mutant strain construction

For clean site-specific mutagenesis, a dual-negative selection-based approach was used as previously described^81^. All strains, primers, and plasmids used for *S*. Tm strain construction are listed in **Table S1**. All plasmid constructs were cloned into *E. coli* DH5α or the DAP-dependent donor strain JKe201^82^. For deletion of *kdgT*, *kdgK*, *edd*, *eda*, *uxuA*, and *idnK*, four primers were used to amplify ∼1 kb upstream and downstream of the given gene and the amplicons were Gibson assembled into the pFOK suicide vector. This construct results in a scarless deletion and the resulting plasmids were confirmed via Sanger sequencing and then transformed into *E*. *coli* DH5α/JKe201. Plasmids were conjugated into wild-type *S*. Tm SL1344 and selected by kanamycin resistance. Stable deletion from the *S*. Tm SL1344 genome required two consecutive homologous recombination events on 20% sucrose with 0.5 µg/mL anhydrous tetracycline (AHT) plates. Two markers were used for negative selection against merodiploids that still have the plasmid integrated into the backbone, using a tetracycline-inducible *P*_tet_ promoter controlling the expression of *sacB* and *I-sceI*. Surviving colonies contained the clean deletion of a given gene and were sequence verified using the appropriate primers (**Table S1**).

### Replication rate measurements during macrophage infection

Approximately 10^6^ RAW 264.7 cells were seeded into 12-well plates (Fisher scientific #08-772-29) and grown overnight for half a doubling. Wild-type and mutant *S.* Tm strains containing the pFccGi reporter plasmid, which encodes mCherry under constitutive expression and GFP under arabinose-inducible expression, were grown overnight in LB with 2% arabinose and appropriate antibiotics. The next day, cells were washed and infected using the protocol described above. Cells were processed at 2, 6, and 10 h.p.i. for flow cytometry. Cells were washed and harvested in flow wash buffer (2% FBS, 0.1% ethylenediaminetetraacetic acid (EDTA) in PBS), stained with LIVE/DEAD stain for 15 min (Thermo Fisher scientific #L34966), fixed and permeabilized for 15 min using Cytofix/Cytoperm (BD Biosciences #554714), washed in permeabilization/wash buffer (BD Biosciences #554714), and resuspended in flow wash buffer. Cells were counted on a Cytek Aurora flow cytometer and analyzed using FlowJo. Live cells were gated based on positive mCherry signal and fold replication was determined using the ratio of the mean GFP and mean mCherry signal from mCherry-positive cells^69^.

